# Complement-dependent loss of inhibitory synapses on pyramidal neurons following *Toxoplasma gondii* infection

**DOI:** 10.1101/2022.07.29.502023

**Authors:** Gabriela L. Carrillo, Jianmin Su, Mikel L. Cawley, Derek Wei, Simran K. Gill, Ira J. Blader, Michael A. Fox

**Affiliations:** Center for Neurobiology Research, Fralin Biomedical Research Institute at Virginia Tech Carilion, Roanoke, Virginia, 24016, USA; Graduate Program in Translational Biology, Medicine, and Health, Virginia Tech, Blacksburg, Virginia, 24061, USA; School of Neuroscience, College of Science, Virginia Tech, Blacksburg, Virginia, 24061, USA; Department of Psychology, Roanoke College, Salem, Virginia, 24153, USA; NeuroSURF Program, Center for Neurobiology Research, Fralin Biomedical Research Institute at Virginia Tech Carilion, Roanoke, Virginia, 24016, USA; Department of Microbiology and Immunology, University at Buffalo, Buffalo, New York, 14203, USA; Department of Biological Sciences, College of Science, Virginia Tech, Blacksburg, Virginia, 24061, USA; Department of Pediatrics, Virginia Tech Carilion School of Medicine, Roanoke, Virginia, 24016, USA

**Keywords:** Toxoplasma gondii, complement, inhibitory synapse, microglia, pyramidal neuron, parvalbumin interneuron

## Abstract

The apicomplexan parasite *Toxoplasma gondii* has developed mechanisms to establish a central nervous system infection in virtually all warm-blooded animals. Acute *T. gondii* infection can cause neuroinflammation, encephalitis, and seizures. Meanwhile, studies in humans, non-human primates, and rodents have linked chronic *T. gondii* infection with altered behavior and increased risk for neuropsychiatric disorders, including schizophrenia. We previously demonstrated that *T. gondii* infection triggers the loss of perisomatic inhibitory synapses, an important source of inhibition on excitatory pyramidal cells, and a type of synapse that is disrupted in neurological and neuropsychiatric disorders. Similar to other instances of inflammation and neurodegeneration, we showed that phagocytic cells (including microglia and infiltrating monocytes) contribute to the loss of these inhibitory synapses. However, in the case of *T. gondii*-induced synapse loss, phagocytic cells target and ensheath the cell bodies of telencephalic neurons. Here, we show that these phagocytic cells specifically ensheath excitatory pyramidal neurons, leading to the preferential loss of perisomatic synapses on these neurons. In contrast, inhibitory cortical interneuron subtypes are not extensively ensheathed by phagocytic cells following infection. Moreover, we show that infection induces expression of complement C3 protein by these excitatory neurons and that C3 is required for the loss of perisomatic inhibitory synapses, albeit not through activation of the classical complement pathway. Together, these findings provide evidence that *T. gondii* infection induces changes in excitatory pyramidal neurons that trigger selective removal of inhibitory perisomatic synapses in the infected neocortex and provide a novel role for complement in remodeling of inhibitory circuits in the infected brain.

## INTRODUCTION

The central nervous system (CNS) is protected by a vascular blood-brain barrier (BBB) that prevents pathogens from entering the brain. However, in some cases, select pathogens have evolved mechanisms to traverse this barrier and invade the CNS. The apicomplexan parasite, *Toxoplasma gondii,* is one such pathogen that enters the brain and can establish a long-lasting CNS infection in almost all warm-blooded animals, including humans (Dubey, 2009, Montoya and Liesenfield, 2004). In fact, it is estimated that over 30% of the global human population are chronically infected with *Toxoplasma* (Flegr et al., 2014, Pappas, Roussos, and Falagas, 2009). Although there are several ways in which humans come into contact with *T. gondii,* we most commonly become infected by ingestion of raw or undercooked meat contaminated by *T. gondii* tissue cysts or by consumption of vegetables and water that have *T. gondii* oocysts shed from the feline definitive host (Hajimohammadi et al., 2022, Jones and Dubey, 2005, Dabritz et al., 2007). Upon oral ingestion, *T. gondii* cysts and oocysts rupture and transform into tachyzoites, the form of the parasite that rapidly grows and disseminates throughout the host (Montoya and Liesenfield, 2004).

This acute phase of the infection is characterized by a severe inflammatory response aimed at controlling the infection, and the tachyzoite form of the parasite responds by differentiating into bradyzoites and forming intracellular cysts. These cysts can persist for the host’s lifetime and they can reactivate. If reactivation occurs in an immunocompetent individual the tachyzoites that emerge will activate a cellular immune response that rapidly and efficiently controls the acute infection. In the absence of such an immune response, or if the immune response is too robust, disease will ensue. In such cases, the brain is the most commonly affected tissue, with patients developing life-threatening seizures and other neurological sequelae (Wong and Remington, 1993, Suzuki and Remington, 1993, Montoya and Liesenfeld, 2004).

Although tachyzoites can invade any nucleated cell, in the brain, they preferentially infect neurons where they transform into cyst-encased bradyzoites that are protected from immune-clearance. (Dubey, 2009, Melzer et al., 2010, Cabral et al., 2016, McConkey et al., 2013, Sims et al., 1989). A substantial body of evidence suggests that chronic *T. gondii* infection alters behaviors in infected hosts (Berdoy et al., 2000, Beste et al., 2014, Vyas et al., 2007). Moreover,although it is not yet understood why, dozens of studies, including a large-scale retrospective study of over 80,000 individuals, indicate a strong association between *T. gondii* infection and an increased risk for developing neuropsychiatric disorders, including schizophrenia (Burgdorf et al., 2019, Dickerson et al., 2014, 2017, 2018, Kano et al., 2020, Wang et al., 2019, Xiao et al., 2018).

Animal models of *T. gondii* infection highlight the impact that this parasite has on brain function by demonstrating that infection leads to seizure development (Brooks et al., 2015, David et al., 2016). In these models, *T. gondii* infection also leads to changes in neurotransmitter synthesis and release (Alsaady et al., 2019, Gatkowska et al., 2016, Martin et al., 2015, Skallova et al., 2006) and altered neural circuits (Brooks et al., 2015, Ihara et al., 2016, Lang et al., Parlog et al., 2015). This is especially true for the inhibitory neurotransmitter, gamma-aminobutyric acid (GABA), and GABAergic circuits in the telencephalon (including the hippocampus and cerebral cortex). For example, studies show *T. gondii* is able to modulate the synthesis of GABA by upregulating expression of GABA synthesis enzymes, GAD65 and GAD67, and downregulating GABA-T, the enzyme responsible for breaking down GABA (Fuks et al., 2012, Bhandage et al., 2020). Additionally, we recently demonstrated that *T. gondii* infection leads to the loss of perisomatic GABAergic synapses from hippocampal and cortical neurons (Carrillo et al., 2020).

The loss of these inhibitory perisomatic synapses following *T. gondii* infection involves the activation of phagocytic cells, which in the infected brain, include both activated microglia and macrophages derived from infiltrating monocytes. We recently showed that these phagocytic cells ensheath neuronal somata and subsequently phagocytose perisomatic inhibitory synapses in the *T. gondii*-infected neocortex (Carrillo et al., 2020). Yet, whether perisomatic inhibitory synapse loss is an indiscriminate process or specific to cell types, and the mechanism underlying synapse loss has remained unclear. Here we sought to address these two important questions. We show that these phagocytic cells preferentially ensheath the somata of excitatory pyramidal neurons in neocortex and that these excitatory neurons differentially express complement C3 in *T. gondii*-infected brains. Moreover, conditional deletion of C3, but not C1qa, rescued the loss of perisomatic inhibitory synapses following infection and substantially reduced neuronal ensheathment, but had no significant effect on the initial phagocyte targeting of these neurons. Thus, our findings demonstrate a novel role for complement in inducing cell-type specific ensheathment and phagocytosis of GABAergic perisomatic synapses in *T. gondii* CNS infection.

## RESULTS

### Differential ensheathment of cortical neurons by IBA1^+^ phagocytes in parasite-infected neocortex

We previously discovered that long-term infection with type II ME49 *Toxoplasma gondii* (> 30 days post infection; Fig 1A) results in the development of seizures and the loss of perisomatic inhibitory synapses in several regions of the mouse telencephalon, including in stratum pyramidalis of the hippocampus and layer V of the cerebral cortex (Carrillo et al., 2020, Brooks et al., 2015). The loss of perisomatic synapses co-occurred with the ensheathment of neuronal cell bodies by IBA1^+^ phagocytic cells (which likely includes both resident microglia and macrophages-derived from peripheral monocytes that infiltrate the brain during infection (Carrillo et al., 2020). The ensheathment of neuronal somata by IBA1^+^ cells in the telencephalon of infected mice is dramatic and noteworthy for at least 3 reasons: 1) neuronal ensheathment and perisomatic synapse loss was widespread throughout the telencephalon at 4 weeks post-infection; 2) some neurons appeared to be almost entirely ensheathed by IBA1^+^ cells; and 3) not all neurons appeared to be ensheathed (Figure 1B). This led to a number of questions: *When do phagocytic cells ensheath neurons following infection? What types of neurons are ensheathed? And what mechanisms drive neuronal ensheathment and perisomatic synapse loss?* Here, we have addressed these important questions.

**Figure 1.**
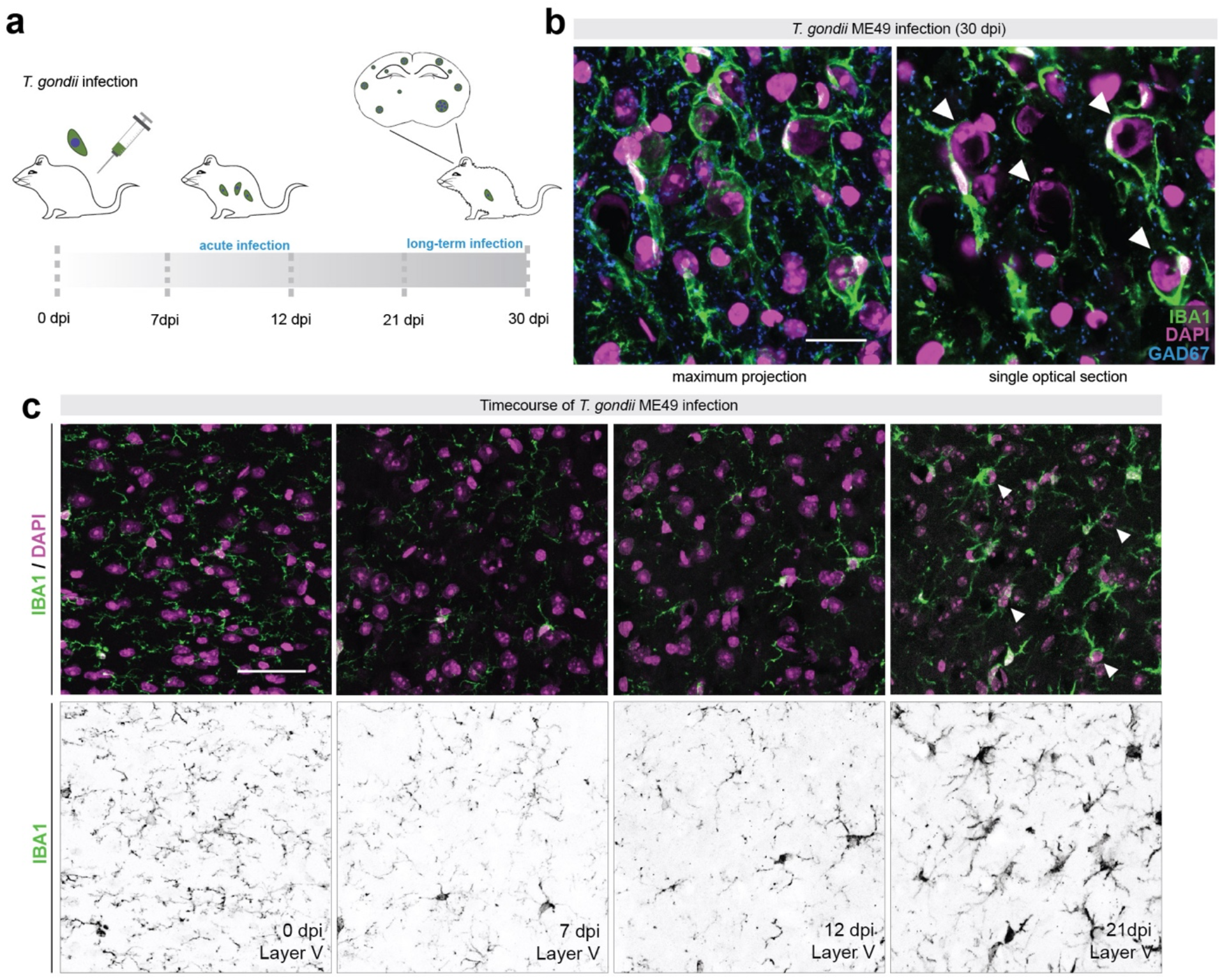
Inhibitory synapse loss and microglial targeting of neuronal somata begins weeks after infection. (a) Schematic representation of *T. gondii* ME49 infection following intraperitoneal injection in adult C57BL/6J mice. (b) Immunostaining of IBA1^+^ phagocytes and GAD67^+^ inhibitory synapses in layer V of neocortex of ME49-infected (30 dpi) mice. Single optical section is shown to highlight cells ensheathed by IBA1^+^ cells (arrows). (c) IHC for IBA1 and DAPI in layer V of cortex at 0 dpi, 7 dpi, 12 dpi, and 21 dpi infection. Arrows depict ensheathed cells at 21 dpi. Scale bars in B: 20 μm and in C: 40 μm

First, we addressed the timing of neuronal ensheathment by phagocytic cells by assessing the distribution of IBA1^+^ cells in the neocortex each week after infection. We show that phagocytes begin to target neurons between 12 and 21 days post infection (dpi), with significant ensheathment of neuronal somata observed at 21 dpi (Figure 1C).

Next, we sought to address whether there was any specificity in terms of which cell types were targeted and ensheathed by IBA1^+^ phagocytes following infection. It is noteworthy that in layer V of neocortex of infected mice, the neurons ensheathed by IBA1^+^ cells appeared pyramidal in morphology (Figure 1B), suggesting excitatory neurons may be preferentially targeted by these phagocytes. To determine if this was indeed the case, brains from infected mice were immunostained for IBA1 and a number of cell-type specific markers allowing us to differentiate between types of excitatory and inhibitory neurons. In layer V of neocortex (where we focus all of these studies), we show that COUP TF1-interacting protein 2^+^ (CTIP2) expressing excitatory pyramidal neurons are extensively ensheathed by phagocytes following infection (Figure 2A-C; Arlotta et al., 2005, Lodato et al., 2011). We measured the extent of the somal surface of these neurons ensheathed by IBA1^+^ cells and found, on average, the surface of pyramidal neurons was over 40% covered by IBA1^+^ cells (Figure 2B, C). Next, we used immunostaining and genetic reporter lines to assess ensheathment of GABAergic interneurons. A subset of somatostatin^+^ (SST), calbindin^+^ (CALB), and parvalbumin^+^ (PV) GABAergic interneurons were contacted by phagocytes in *Toxoplasma*-infected cortex, although this is also the case in mock-infected brains. While all 3 types of interneurons exhibited higher percentage of ensheathment in infected brains than in mock-infected brains (Figure 2D-L), the level of ensheathment remained significantly lower (~15% ensheathed per cell) compared to pyramidal neurons (Figure 2). Thus, it appears that excitatory neurons are preferentially targeted and ensheathed by IBA1^+^ phagocytes following long-term infection with *Toxoplasma gondii*.

**Figure 2.**
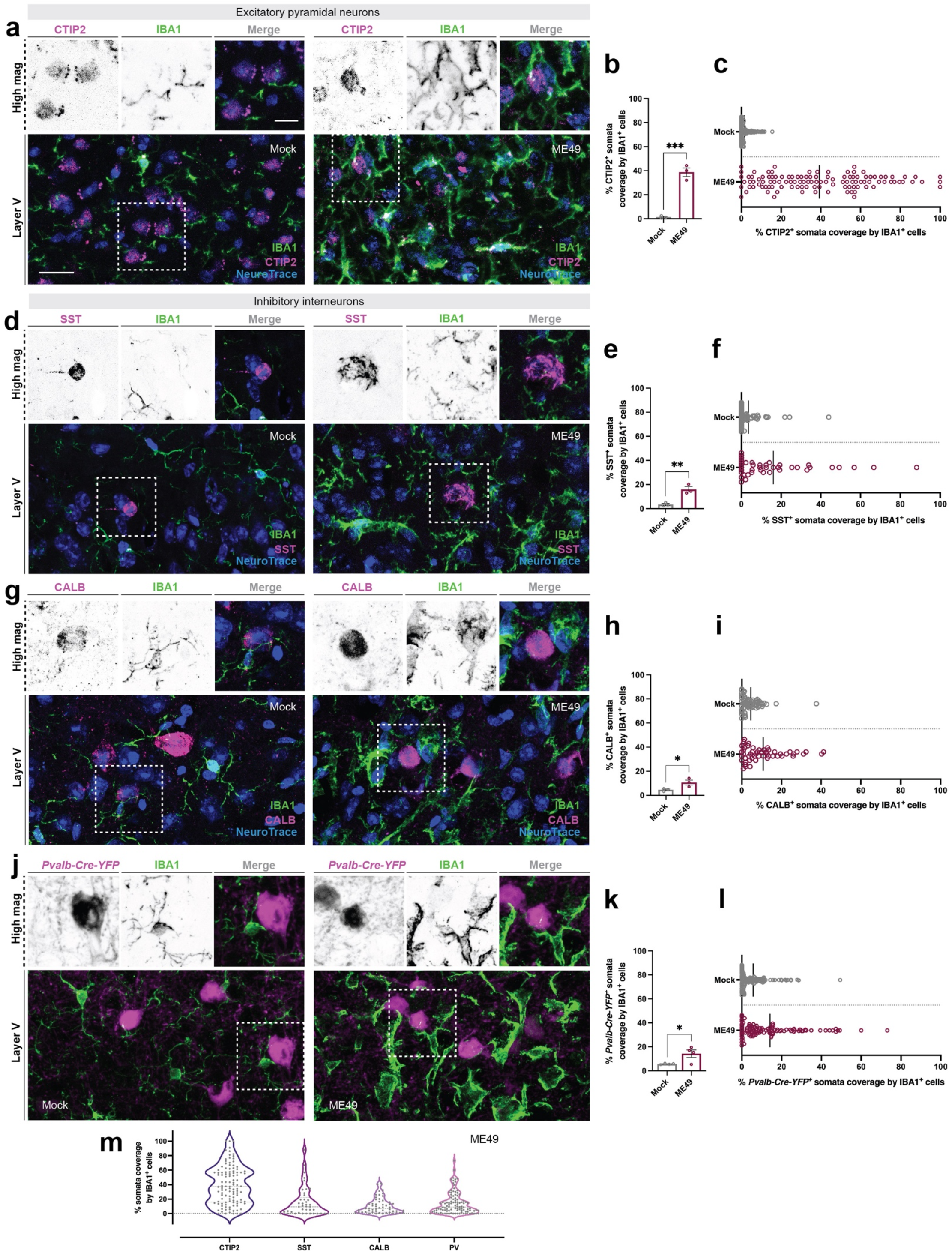
Excitatory pyramidal neurons are preferentially ensheathed by microglia. (a) IHC for layer V neurons (CTIP2), NeuroTrace and IBA1 in layer V of neocortex at 30 dpi (ME49) compared to mock-infection (Mock). (b) Quantification of CTIP2^+^ NeuroTrace^+^ soma coverage by IBA1^+^ cells in mock- and ME49-infected neocortex. Each data point represents the average of one biological replicate and bars depict mean ± SEM. Asterisks (***) indicate *P* < 0.001 by Student’s *t*-test (n=3 mice per condition). (c) Distribution plot of individual CTIP2^+^ NeuroTrace^+^ somata analyzed and pooled from biological replicates in (B). Bar represents mean percent somata coverage by all IBA1^+^ cells. (d) IHC for SST, NeuroTrace, and IBA1 in layer V of neocortex (ME49) compared to mock-infection (Mock). (e) Quantification of SST^+^ NeuroTrace^+^ soma coverage by IBA1^+^ cells in mock- and ME49-infected neocortex. Each data point represents the average of one biological replicate and bars depict mean ± SEM. Asterisks (**) indicate *P* < 0.01 by Student’s *t*-test (n=3 mice per condition). (f) Distribution plot of individual SST^+^ NeuroTrace^+^ somata analyzed and pooled from biological replicates in (E). Bar represents mean percent somata coverage by all IBA1^+^ cells. (g) IHC for CALB, NeuroTrace, and IBA1 in layer V of neocortex (ME49) compared to mock-infection (Mock). (h) Quantification of CALB^+^ NeuroTrace^+^ soma coverage by IBA1^+^ cells in mock- and ME49-infected neocortex. Each data point represents the average of one biological replicate and bars depict mean ± SEM. Asterisks (*) indicate *P* < 0.05 by Student’s *t*-test (n=3 mice per condition). (i) Distribution plot of individual CALB^+^ NeuroTrace^+^ somata analyzed and pooled from biological replicates in (H). Bar represents mean percent somata coverage by all IBA1^+^ cells. (j) IHC for IBA1 in *Pvalb-Cre-YFP* shows minimal ensheathment of PV^+^ inhibitory interneurons in layer V of neocortex at 30 dpi (ME49) compared to mock-infection (Mock). (k) Quantification of *Pvalb-Cre-YFP*^+^ soma coverage by IBA1^+^ cells in mock- and ME49-infected neocortex. Each data point represents the average of one biological replicate and bars depict mean ± SEM. Asterisks (*) indicate *P* < 0.05 by Student’s *t*-test (n=4 mice per condition). (l) Distribution plot of individual *Pvalb-Cre-YFP*^+^ somata analyzed and pooled from biological replicates in (K). Bar represents mean percent somata coverage by all IBA1^+^ cells. (m) Violin-plot showing the distribution of ensheathment by IBA1^+^ phagocytes (as a % of somal coverage) for 4 types of neocortical neurons (Data from c, f, i, l). Scale bars in A, D, G, J: 20 μm and in A, D, G, J high magnification images: 10 μm

It is possible that certain cell types, such as GABAergic interneurons, are preferentially lost following infection, ensheathment, and synapse loss. This could lead to lower numbers of highly ensheathed GABAergic interneurons in our counts (and to alterations in GAD67 distribution (Brooks et al., 2015)). To test this possibility, we used riboprobes against *Gad1* and *Vglut1* to label inhibitory and excitatory neurons, respectively, by *in situ* hybridization (ISH). In line with our previous assessment of neuronal loss in the infected neocortex (Carrillo et al., 2020), we did not observe significant changes in the number of excitatory or inhibitory cells in layer V at 30 days post infection (Figure S1).

### Perisomatic inhibitory synapses are not lost from the somata of PV^+^ interneurons follow infection

We interpret the above results to suggest that pyramidal neurons most likely express or release a factor that attracts phagocytes to ensheath their somata. However, the somata of these neurons are studded with perisomatic synapses, so an alternative possibility is that phagocytes are instead attracted to these synapses following infection. In layer V, the majority of perisomatic synapses on excitatory pyramidal cells are formed by fast-spiking, PV-expressing interneurons. Importantly, however, PV^+^ cells also provide substantial perisomatic inhibition onto other PV^+^ cells (Figure 3A-B). Since PV^+^ interneurons are not dramatically ensheathed by IBA1^+^ cells following infection (Figure 2J-L), this suggests that perisomatic nerve terminals are not attracting IBA1^+^ cells to ensheath neuronal somata and phagocytose these nerve terminals following infection. To directly test the latter possibility, we took advantage of our genetic reporter line labeling PV cells to probe whether perisomatic synapse loss also occurred on these inhibitory PV^+^ interneurons. We observed no significant decrease in the number of GABAergic perisomatic synapses on PV^+^ interneurons (labeled by immunostaining for glutamic acid decarboxylase 67; GAD67) between mock- and ME49-infected brains (Figure 3C-E). Additionally, we quantified the number of PV^+^ interneurons in layer V and in the neighboring cortical layers (IV-VI) and found no change in PV^+^ cell numbers following infection (Figure 3F-G). Together, these results show that PV^+^ cells are not extensively ensheathed by IBA1^+^ phagocytes following infection, nor do they lose perisomatic inhibitory inputs form their somata. This suggests that post-synaptic signals arising from excitatory pyramidal cells are likely serving as cues for microglia to ensheath excitatory somata and remove inhibitory inputs.

**Figure 3.**
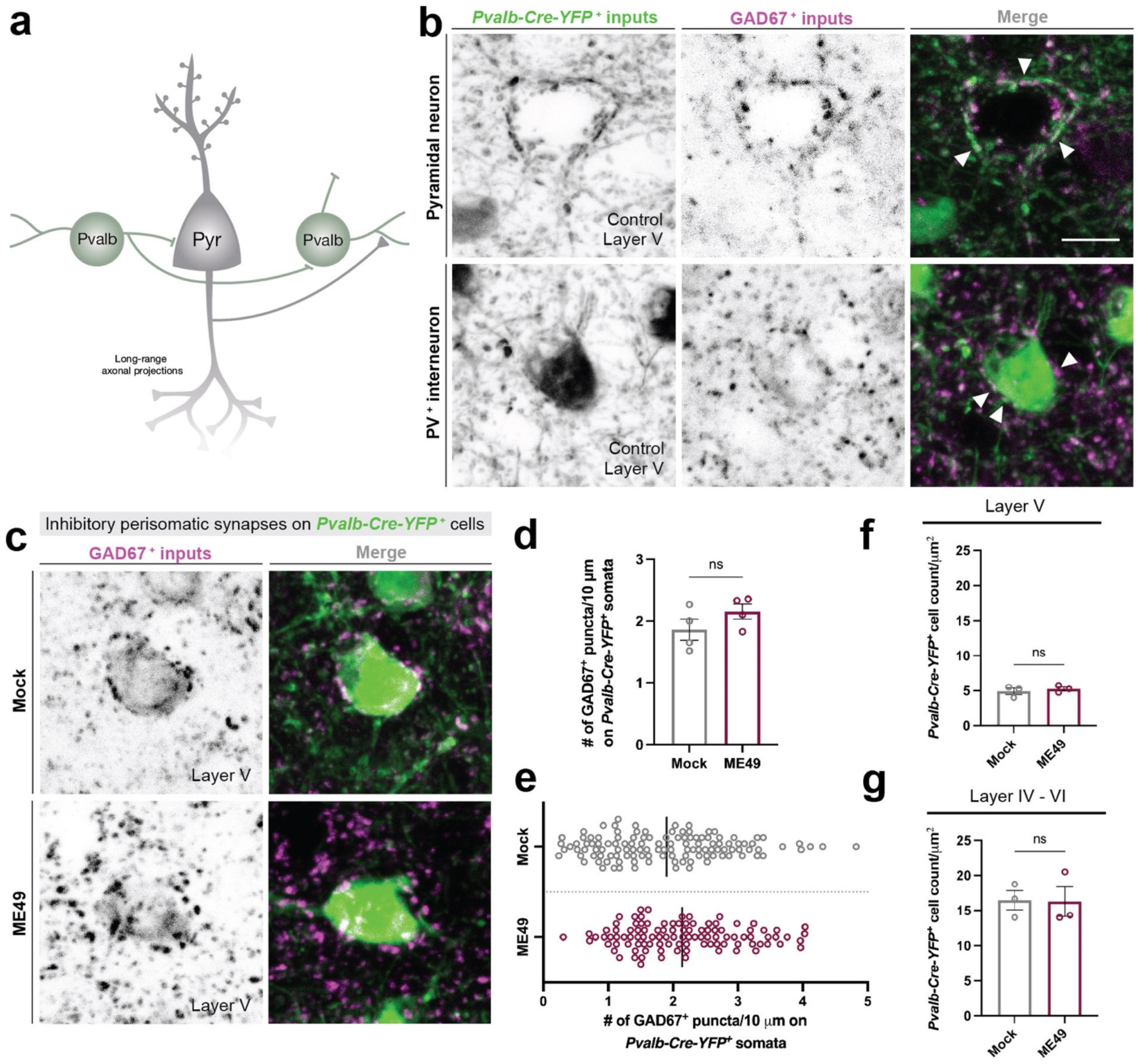
Inhibitory perisomatic synapses are not lost on PV^+^ interneurons. (a) Schematic illustrating PV inhibitory interneuron and excitatory pyramidal connectivity in cortical layer V. (b) IHC for GAD67 in *Pvalb-Cre-YFP* mice shows *YFP+* inputs on both pyramidal neurons (based on morphology) and on PV interneurons. White arrowheads indicate inhibitory perisomatic synapses. (c) IHC for GAD67 in mock- and ME49-infected (30 dpi) *Pvalb-Cre-YFP* mice. (d) Quantification of GAD67^+^ synapses on *YFP*^+^ somata in mock- and ME49-infected Pvalb-Cre-YFP mice. Each data point represents one biological replicate and bars depict mean ± SEM. No significant difference (ns) indicates *P* > 0.05 by Student’s *t*-test (n=4 mice per condition). (e) Distribution plot of *YFP^+^* somata analyzed and pooled from biological replicates in (D). Bar represents mean number of GAD67^+^ synapses per *Pvalb-Cre-YFP*^+^ somata in mock- or ME49-infected neocortex. (f) Quantification of *Pvalb-Cre-YFP*^+^ cell count in layer V of mock- and ME49-infected cortex. Each data point represents one biological replicate and bars depict mean ± SEM. No significant difference (ns) indicates *P* > 0.05 by Student’s *t*-test (n=3 mice per condition). (g) Quantification of *Pvalb-Cre-YFP*^+^ cell count in layer IV - VI of mock- and ME49-infected cortex. Each data point represents one biological replicate and bars depict mean ± SEM. No significant difference (ns) indicates *P* > 0.05 by Student’s *t*-test (n=3 mice per condition). Scale bars in B and C: 10 μm

### Differential expression of complement components following *T. gondii* infection

We next sought to identify what some of the cues might be that are necessary for IBA1^+^ phagocyte ensheathment of excitatory pyramidal neurons and the targeted loss of perisomatic synapse on these neurons. Activation of the classical complement pathway has been shown to control synaptic pruning during developmental refinement as well as in the neurodegenerating brain (Stevens et al., 2007, Cong et al., 2021, Alawieh et al., 2021, Werneburg et al., 2020, Hammond et al., 2020). Moreover, several studies report upregulation of complement components in the *Toxoplasma*-infected brain (Shinjyo et al., 2020, Huant et al., 2018, Xiao et al., 2016). We similarly observed significant increase of classical components *C1qa* and *C3* in infected cortex by RT-qPCR *(C1qa:* mock = 1 ± 1.3 fold-change; ME49 = 6.5 ± 4.2; *C3*: mock = 1 ± .95; ME49 = 267 ± 13.5). Based on this, we hypothesized that these complement pathway components might be involved. In the inflamed or injured brain, microglia provide a major source of *C1qa* mRNA, however it remains unknown which cells express *C1qa* and *C3* in the *Toxoplasma*-infected brain. To determine the source of complement components in the *T. gondii*-infected brain, we generated riboprobes against *C1qa* and *C3* and performed ISH. In the infected brain, *C1qa* mRNA expression patterns resembled phagocyte morphology. Immunolabeling for IBA1 following *in situ* hybridization confirmed the presence of *C1qa* mRNA within IBA1^+^ cells in both mock- and ME49-infected brains. In fact, the majority of IBA1^+^ cells in both sets of brains expressed *C1qa* although expression appeared higher in the infected brain (Figure 4A-C). Additionally, a population of cells expressing *C1qa* that are not labeled with IBA1 are present (Figure S3A). ISH against *C3* mRNA similarly revealed expression by IBA1^+^ phagocytes in the infected brain. However, in contrast to *C1qa* mRNA, *C3* mRNA expression was not detectable in the mock-infected brain.

**Figure 4.**
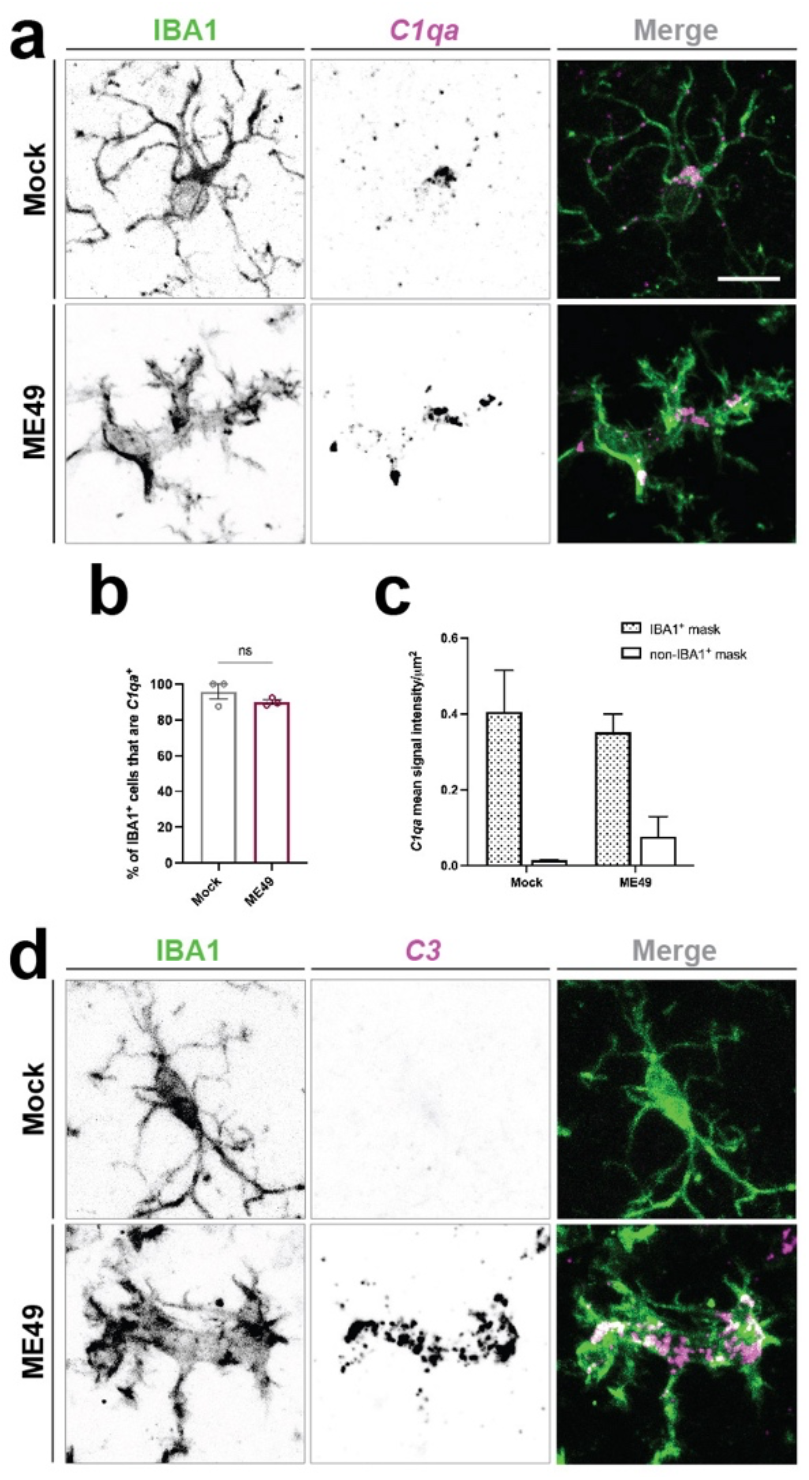
Complement mRNA is upregulated by immune cells in the infected brain. (a) ISH for *C1qa* and IHC for IBA1 in layer V of neocortex of mock- and ME49-infected mice. (b) Quantification of IBA1^+^ cells expressing *C1qa* mRNA in mock- and ME49-infected cortex. Each data point represents one biological replicate and bars depict mean ± SEM. No significant difference (ns) indicates *P* > 0.05 by Student’s *t*-test (n=3 mice per condition). (c) Quantification of *C1qa* mRNA signal intensity within IBA1^+^ cells and outside of IBA1^+^ cells. Each data point represents one biological replicate and bars depict ± SEM. (d) ISH for *C3* and IHC for IBA1 in layer V of neocortex of mock- and ME49-infected mice. Scale bars in A and D: 10 μm

We next sought to examine neuronal subtype expression for *C1qa* and *C3* in mock- and ME49-infected brains. To accomplish this, we performed *in situ* hybridization for *C1qa* and *C3* and genes expressed by all neurons (Synaptotagmin 1; *Syt1),* GABAergic neurons (Glutamic acid decarboxylase 1; *Gad1)* or layer V pyramidal neurons (Nephrenectin; *Npnt)* (Figure S2; Su et al., 2021). While we observed instances of *C1qa* expression by non-IBA1^+^ cells, we failed to observe any *C1qa* expression by *Syt1*^+^ neurons (Figure S3). However, we did find several instances of *C3* mRNA expression by both *Syt1^+^* and *Npnt^+^* neurons in the ME49-infected cortex (Figure 5A, B). Importantly, we did not observe *C3* expression by *Gad1*^+^ inhibitory cells (Fig 5C).

**Figure 5.**
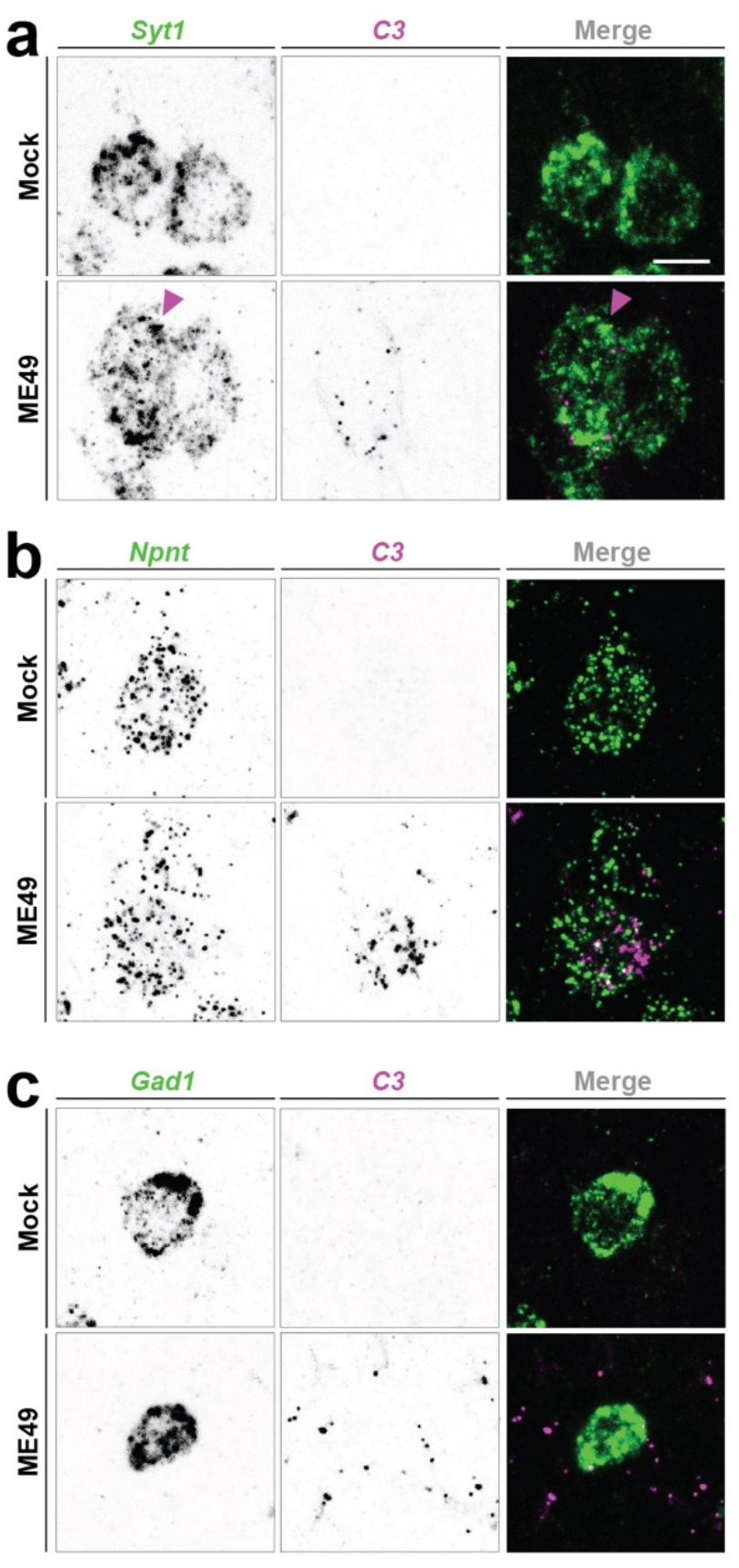
C3 is expressed by some excitatory pyramidal neurons following *T. gondii* infection. (a) ISH for *Syt1* and *C3* in mock- and ME49-infected mice. Neuronal expression of *C3* mRNA was detected in some neurons in ME49-infection. (b) ISH for *Npnt* and *C3* in mock- and ME49-infected mice. *C3* mRNA was detected in some *Npnt-*expressing pyramidal neurons in layer V of neocortex. (c) ISH for *Gad1* and *C3* show no expression of *C3* mRNA by inhibitory neurons in ME49-infection. Scale bars in A, B and C: 10 μm

### C3 is required for inhibitory perisomatic synapse loss in a non-classical pathway-dependent manner

The select expression of C3 by some pyramidal neurons, but not interneurons, suggests that the complement pathway may contribute to neuronal ensheathment by IBA1^+^ phagocytes and perisomatic synapse loss. To test this, we assessed infection-induced IBA1^+^ ensheathment of neurons and perisomatic synapse loss in mice lacking C3. Following most types of infection or injury, resident microglia quickly expand in the brain, and monocytes from the periphery are recruited into the brain to aid with pathogen and debris clearance (D’Mello et al., 2009). We observe this same process following *T. gondii* infection in WT mice (Fig 6A, B). Interestingly, in mice that lack C3 globally, we also see an increase in the number of phagocytes following infection, however, the number is slightly but significantly less than in WT mice (Figure 6 A,B). We failed to observe significant differences in the percentage of IBA1^+^ phagocytes that contact neurons in WT and C3^-/-^ infected brains. However, we observed a significant decrease in neuronal ensheathment by IBA1^+^ cells (Figure 6C-F). In fact, loss of C3 led to ensheathment levels that quantitatively and qualitatively resembled what levels we observed on inhibitory interneurons in WT infected mice (compares Figure 2 and 6).

**Figure 6.**
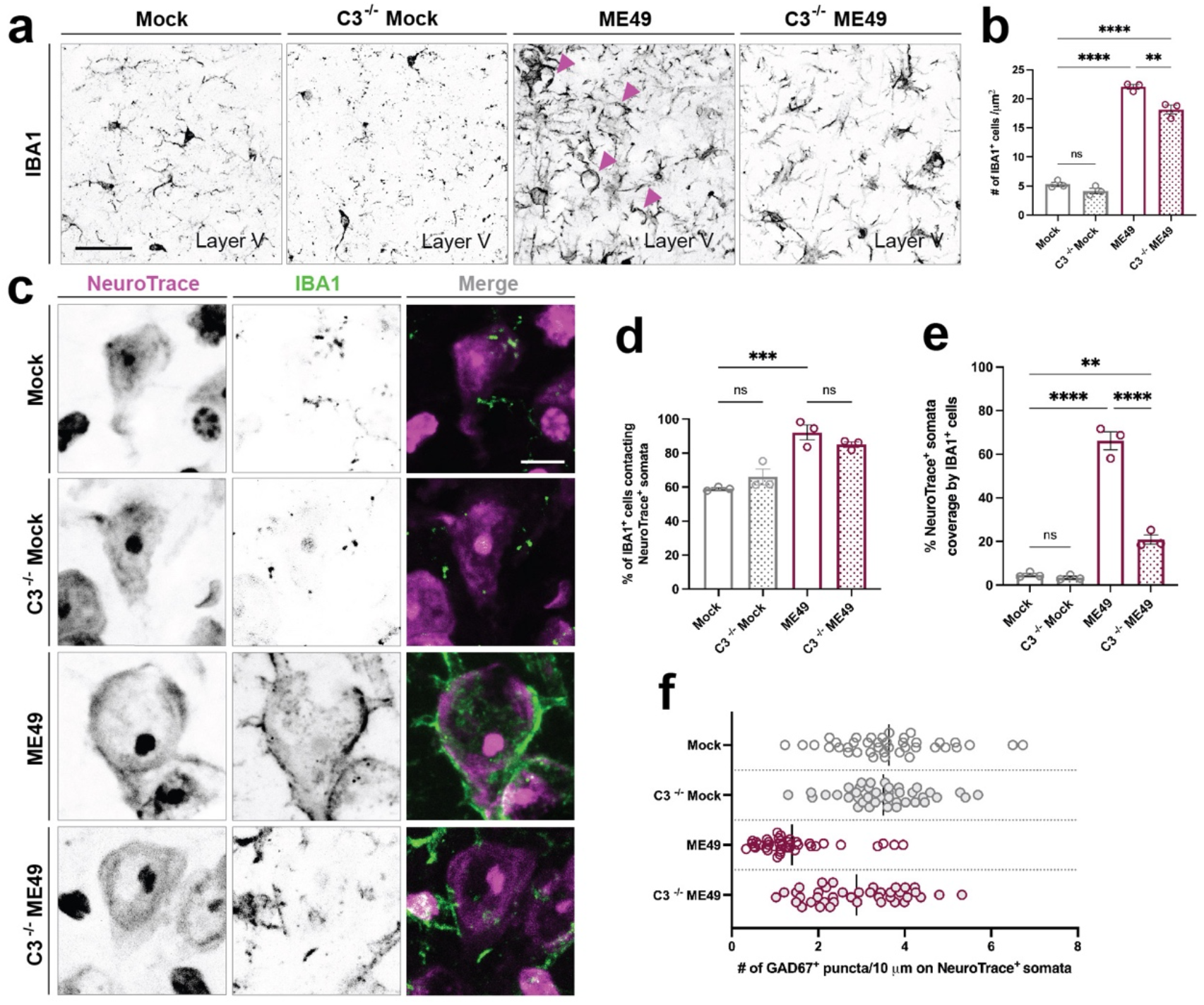
Reduced neuronal ensheathment by IBA1^+^ phagocytes in the absence of complement C3. (a) IHC for IBA1 in layer V of neocortex of C3^-/-^ and littermate control mice mock-infected and infected with ME49 parasites. (b) Quantification of the number of IBA1^+^ cells in layer V of neocortex in C3^+/+^ mock-, C3^-/-^ mock, C3^+/+^ ME49-, and C3^-/-^ ME49-infected brains. Asterisks (**) indicate *P* < 0.01, (****) indicate *P* < 0.0001 by one-way ANOVA with Sidak’s multiple comparison correction (n=3 mice per condition). (c) IHC for IBA1 and NeuroTrace in layer V of neocortex of C3^+/+^ mock-, C3^-/-^ mock, C3^+/+^ ME49- and C3^-/-^ ME49-infected mice. (d) Quantification of the percentage of IBA1^+^ cells contacting NeuroTrace^+^ somata in cortical layer V in in C3^+/+^ mock-, C3^-/-^ mock, C3^+/+^ ME49-. and C3^-/-^ ME49-infected brains. Each data point represents one biological replicate and bars depict mean ± SEM. Asterisks (***) indicate *P* < 0.001 and (ns) indicates *P* > 0.05 by one-way ANOVA with Sidak’s multiple comparison correction (n=3 mice per condition). (e) Quantification of the percentage of NeuroTrace^+^ soma coverage by IBA1^+^ cells in cortical layer V in C3^+/+^ mock-, C3^-/-^ mock, C3^+/+^ ME49-. and C3^-/-^ ME49-infected brains. Each data point represents one biological replicate and bars depict mean ± SEM. Asterisks (**) indicate *P* < 0.01, (****) indicate *P* < 0.0001 by one-way ANOVA with Sidak’s multiple comparison correction (n=3 mice per condition). (f) Distribution plot of individual NeuroTrace^+^ somata analyzed and pooled from biological replicates in (D). Bar represents mean percent somata coverage by IBA1^+^ cells. Scale bars in A: 40 μm and in C: 10 μm

Next, to determine if the reduced ensheathment of neurons in the absence of C3 resulted in less phagocytosis and loss of synapses following infection, we assessed the distribution of CD68 (a marker of phagosomes) and of inhibitory perisomatic synapses (Carrillo et al., 2020). In C3 ^-/-^ infected mutants, we saw a decrease in the distribution of CD68 immunoreactivity within layer V (Figure 7A-B). Importantly, we found that not all phagocytes showed a loss or reduction in CD68 expression. Areas of the cortex where phagocytes accumulated (perhaps denoting regions of monocyte infiltration or macrophage response to tachyzoites within the brain parenchyma), for example, still expressed high levels of CD68, suggesting that the increase in phagocytic activity in ensheathing phagocytes (but not all phagocytes) was dependent on C3 activation (Figure 7C). We next assessed the number of GABAergic inhibitory synapses and saw no significant loss of perisomatic synapses in the neocortex of infected C3 ^-/-^ mutants (Figure 7D-F). Since the number of perisomatic synapses in C3 ^-/-^ mock-infected cortex was unchanged from WT mock-infected cortex, we conclude that these findings demonstrate C3 is required for the phagocytosis of inhibitory perisomatic synapses in *T. gondii* infection.

**Figure 7.**
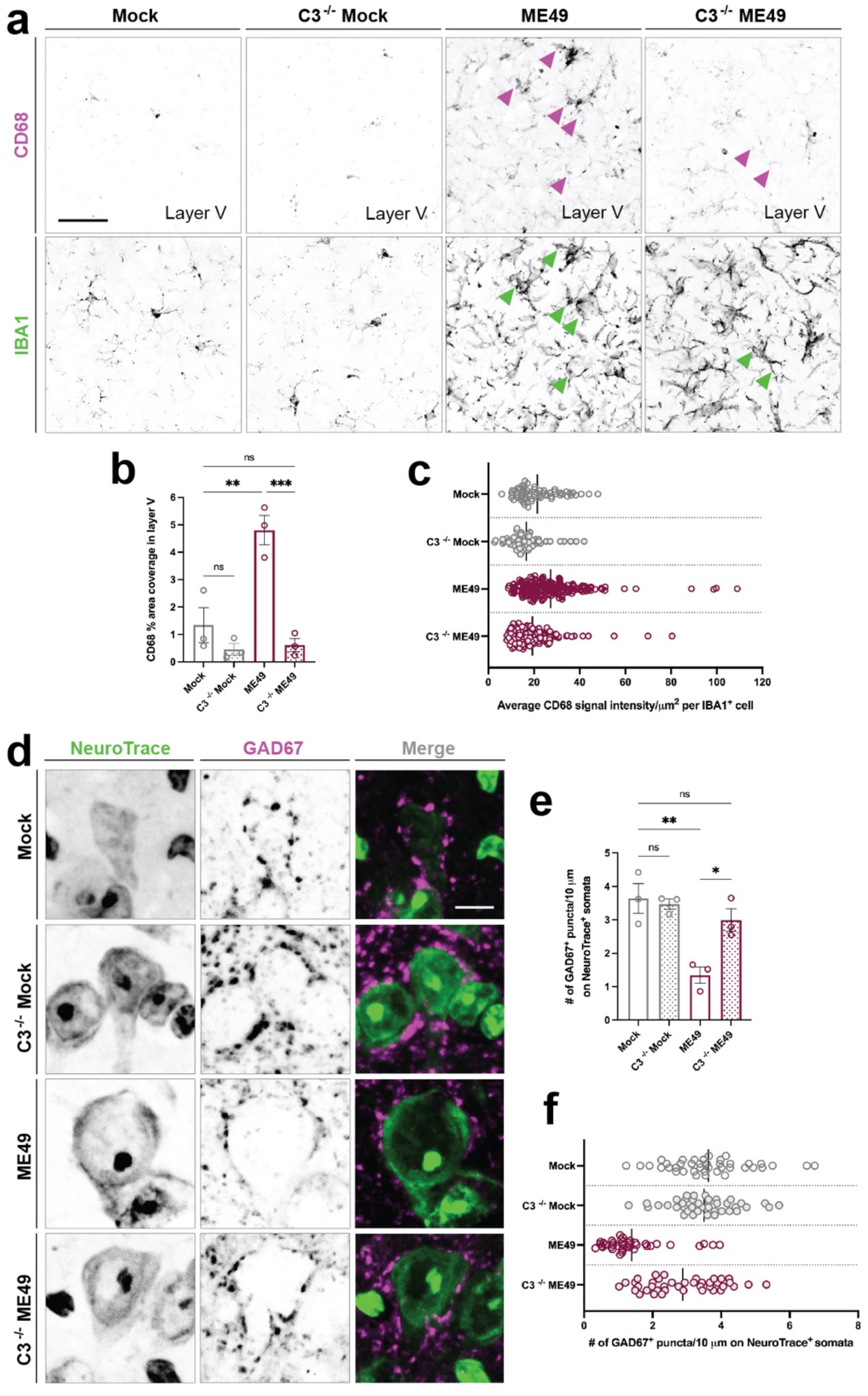
C3 is required for phagocytosis of inhibitory perisomatic synapses. (a) IHC for IBA1 and CD68 show decreased expression by IBA1^+^ cells in layer V of neocortex in C3^-/-^ mice following *T. gondii* infection compared to C3^+/+^ ME49-infected mice. Magenta arrowheads point to CD68 expression with IBA1^+^ cells. Green arrowheads point to IBA1^+^ cells (b) Quantification of CD68 immunoreactivity percentage area coverage in cortical layer V in C3^+/+^ mock-, C3^-/-^ mock, C3^+/+^ ME49-. and C3^-/-^ ME49-infected brains. Each data point represents one biological replicate and bars depict mean ± SEM. Asterisks (**) indicate *P* < 0.01, (***) indicate *P* < 0.001, and (ns) *P* > 0.05 by one-way ANOVA with Sidak’s multiple comparison correction (n=3 mice per condition). (c) Distribution plot of individual IBA1^*+*^ cells analyzed and pooled from images in (B). Bar represents mean number of CD68 signal intensity within IBA1^+^ cells. (d) IHC for GAD67 and NeuroTrace in layer V of neocortex of C3^+/+^ mock-, C3^-/-^ mock, C3^+/+^ ME49-. and C3^-/-^ ME49-infected mice. (e) Quantification of GAD67^+^ synapses on NeuroTrace^+^ somata in cortical layer V in C3^+/+^ mock-, C3^-/-^ mock, C3^+/+^ ME49-. and C3^-/-^ ME49-infected brains. Each data point represents one biological replicate and bars depict mean ± SEM. Asterisks (*) indicate *P* < 0.05, (**) indicate *P* < 0.01, and (ns) *P* > 0.05 by one-way ANOVA with Sidak’s multiple comparison correction (n=3 mice per condition). (f) Distribution plot of individual NeuroTrace^*+*^ cells analyzed and pooled from images in (E). Bar represents mean number of GAD67^+^ synapses on NeuroTrace^+^ somata. Scale bars in A: 40 μm and in D: 10 μm

After establishing that *T. gondii* infection-induced synapse loss is C3-dependent, we sought to determine if this was mediated through activation of the classical pathway. A number of studies have demonstrated that synapse loss in the diseased or inflamed brain involve the classical complement pathway (Hong et al., 2016, Wang et al., 2020, Datta et al., 2020, Carvalho et al., 2019). To test this, we used targeted mouse mutants that lack C1qa, as enzymatic cleavage of this protein activates the classical pathway. We found that removal of C1qa did not prevent *T. gondii-induced* synapse loss (Figure S4A-C). Thus, the classical complement pathway is not involved in the loss of synapses in the *Toxoplasma*-infected brain, similar to what is seen in neurodegeneration (Werneburg et al., 2020).

## DISCUSSION

CNS infection with *Toxoplasma gondii* can lead to behavioral alterations, seizures, and increased risk for the development of several neuropsychiatric disorders, some, or all, of which may arise from dysfunction in GABAergic neurotransmission and circuitry. Data from our previous studies demonstrate that infection by type II ME49 *T. gondii* leads to the loss of GABAergic perisomatic synapses and the acquisition of spontaneous seizures (Brooks et al., 2015, Carrillo et al., 2020). Dysfunction or perturbation of these inhibitory perisomatic synapses have been linked to both seizures and neuropsychiatric disorders, such as schizophrenia (Belforte et al., 2010, Gonzalez-Burgos et al., 2011, Gonzalez-Burgos et al., 2010, Gonzalez-Burgos and Lewis, 2012, Hamm et al., 2017, Lewis et a., 2011, Mukherjee et al., 2019, Schwaller et al., 2004, Wohr et al., 2015). In the case of *T. gondii* infection, activated microglia and/or peripheral monocytes that infiltrate the brain following infection, ensheath neuronal somata and phagocytose a significant number of the GABAergic perisomatic synapses on these cells (Carrillo et al., 2020). Here, we discovered that the loss of GABAergic perisomatic synapses in the *T. gondii*-infected cerebral cortex occurs in a cell-type specific manner in that phagocytic cells preferentially ensheath excitatory pyramidal cells, leading to the loss of inhibitory perisomatic synapses on this cell type, but not on others. We further demonstrate that this process is complement-dependent but is not mediated by the classical pathway. Together, these findings highlight a new role for complement in orchestrating phagocyte cell-type specific remodeling in the *T. gondii*-infected brain.

### Cell-type specific neuron-glia interactions in parasite-infected brain

Microglia are key regulators of neural circuitry with established roles in immunosurveillance and circuit remodeling in the central nervous system. In the developing brain, microglia eliminate excess neural precursor cells and synapses to maintain homeostasis and refine circuitry (Cunningham et al., 2013, Paolicelli et al., 2011, Li et al., 2012, Schafer et al., 2012). Upon injury or disease to the brain, microglia can temporarily or permanently remodel circuitry by employing both pathogenic and protective mechanisms (Chen et al., 2014, Chen et al., 2012, Kerschensteiner et al., 2003, Hellwig et al., 2013). At the initial stages of the infection, *T. gondii* infection follows a similar pattern of widespread microglial activation and increase in phagocytic cells (here, likely attributed to both microglial proliferation and monocyte recruitment from the periphery), as is commonly seen in other, but not all, types of neuroinflammation or neurodegeneration (Borges et al., 2003, Feng et al., 2019, Hagan et al., 2020). Moreover, *T. gondii* infection induces microglia to extensively ensheath the somata of neurons, a process first described in the peripheral central nervous system following injury to the facial nerve (Blinzinger and Kreutzberg, 1968, Shibasaki et al., 2007, Chen et al., 2014, Wan et al., 2020). Interestingly, ensheathment of neurons in models of induced epilepsy has been described as a transient process, whereby microglia temporarily displace perisomatic synapses as a response to aberrant GABAergic signaling from presynaptic nerve terminals, thus serving as a protective mechanism to the neuron (Wan et al., 2020). Furthermore, displacement of GABAergic perisomatic synapses results in an increase in synchronized neuronal activity which triggers neuronal expression of anti-apoptotic and neurotrophic molecules as neuroprotection (Chen et al., 2014). In *T. gondii* infection, however, we find that GABAergic perisomatic synapse loss is not a result of temporary displacement, but rather is a process where presynaptic nerve terminals are removed by phagocytosis. If widespread perisomatic synapse loss continues throughout the course of the infection, it may contribute to neuronal death seen in later stages of chronic *T. gondii* infection (Cabral et al., 2016, Mendez et al., 2021).

In the current study, we discovered *T. gondii* infection triggers microglia to specifically ensheath excitatory pyramidal neurons in neocortex and that GABAergic perisomatic synapse loss occurs on these excitatory cells. These findings are in line with other models of epilepsy that show preferential ensheathment of excitatory cortical neurons and subsequent displacement of GABAergic perisomatic synapses from these cells (Wan et al., 2020). Microglia, however, are not the only ones that preferentially interact with excitatory neurons. Recent studies assessing neurons that are injected with *T. gondii* proteins (but do not contain the parasite itself) shows the parasite, like microglia, also has preference for targeting excitatory neurons (Koshy et al., 2012, Mendez et al., 2021). Thus, excitatory cells could be secreting signaling molecules such as neurotrophins and neuropeptides that attract both microglia and parasites to these cells. Alternatively, phagocytes may be attracted to these cells as a response to parasite-modulation of host cell machinery or neuronal stress within *T. gondii*-protein-injected neurons. A third possibility is that instead of being attracted to excitatory cells, microglia are attracted to the inhibitory perisomatic synapses. Microglia express a variety of receptors for neurotransmitters, including GABA receptors, that allow them to sense neuronal activity (Pocock and Kettenmann, 2007, Krabbe et al., 2012, Seifert et al., 2011, Kuhn et al., 2004). In *T. gondii* infection, microglia upregulate the expression of GABA-A receptors, which aids in their migration, therefore, it is possible that perisomatic synapses could be releasing an increase in GABA that attracts phagocytes to neuronal somata (Bhandage et al., 2019, Bhandage et al., 2020). However, this is unlikely to be the case since perisomatic inhibitory synapses remain on PV inhibitory interneurons following *T. gondii* infection.

### Differential expression of complement as a cue for microglia ensheathment of excitatory neurons leading to perisomatic synapse phagocytosis

An important discovery in these studies is that along with microglia and monocyte expression of complement components, *T. gondii* infection also induces the differential expression of C3 by excitatory pyramidal cells. Neuronal expression of complement is not an entirely novel concept. In animal models of Alzheimer’s, where synapse loss is an early manifestation of the neurodegenerative process via the complement pathway, neuronal expression of complement C1q has been reported and linked to synapse elimination (Selkoe 2002, Bialas and Stevens, 2013, Hong et al., 2016). Similarly, in chronic cases of multiple sclerosis, microglial clusters (although less extensive than we see in *T. gondii* infection), are associated with neuronal production of C3. In these studies, expression of C3 and microglial clusters only occurred after prolonged disease, and was not observed during acute disease (Michailidoi et al., 2016). In our studies, it is important to note that several instances of neuronal C3 production (by excitatory pyramidal cells only) were observed at 30 days after infection, a timepoint of long-term infection where significant perisomatic inhibitory synapse loss already occurs. This raises important questions as to the timing and duration of complement expression by neurons in the *T. gondii* infected brain. *In vivo* assessment of neuronal C3 expression throughout the course of the infection, and targeted downregulation of C3 in neurons, will be important in establishing neuronal C3 as a driver in *T. gondii*-induced synapse loss. Since phagocytes were also observed to upregulate expression of C3 in *T. gondii* infection, the role of phagocyte C3 in inducing synapse loss should also be carefully examined. This raises the possibility that excitatory pyramidal neuron expression of C3 may also be accompanied by inhibitory neuron upregulation of complement regulators as a protective mechanism against microglia secreted C3 (Zhu et al., 2020, Gonzalez, et al., 2021). Of note, *T. gondii* tachyzoites are able to resist complement-mediated killing by recruiting host-derived complement regulatory proteins (Factor H regulator of the alternative pathway and C4b-binding protein of the classical and lectin pathways) to the parasite’s surface and inactivate surface-bound C3 (Fuhrman and Joiner, 1989, Sikorski et al., 2020). It is possible that intracellular *T. gondii* cysts may employ a similar strategy to avoid complement-mediated phagocytosis or lysis of their neuronal host. Overall, our findings suggest that neurons might be playing a more significant role in initiating circuit remodeling in degeneration or prolonged inflammation and should be examined in other models of neurodegeneration and disease.

Our data suggest that C3 is required for neuronal ensheathment and phagocytosis of perisomatic inhibitory synapses, but is not required for the initial targeting of excitatory pyramidal neurons. This was surprising to us based on the role of C3 in chemoattraction (Chen et al., 2021). This begs the question, what are the chemoattractants that initiate selective microglial targeting of excitatory cells in the *T. gondii*-infected brain? Under pathological conditions, activated microglia will migrate towards the site of injury by detecting different classes of membrane-bound and secreted chemoattractants (Fan et al., 2017, Hu et al., 2014, Mazaheri et al., 2017). One example of these is the inhibitory neurotransmitter, GABA. In the developing cortex, secreted GABA will attract GABA_B_ receptive microglia, and subsequently induce a transcriptional synapse remodeling program within these immune cells to trigger synapse phagocytosis (Favuzzi et al., 2021). In *T. gondii* infection, GABA-induced chemotaxis of microglia may account for the selective phagocytosis of inhibitory perisomatic synapses, but seems unlikely to explain selectivity for excitatory pyramidal neurons. The identification of such cell-type specific chemoattractants in *T. gondii* infection will require further investigation.

## Acknowledgements

This work was supported by National Institute of Health grants NS105141, AI124677, and F99NS120596. We thank the LaMantia and Farris labs for generously supplying some antibodies used in these experiments.

## Author contributions

GLC designed the study, performed experiments, analyzed data, and wrote the manuscript. JS performed experiments. MC and DW analyzed data. IJB and MAF contributed conceptually to design of study and edited manuscript.

---

**Supplemental Figure 1.**
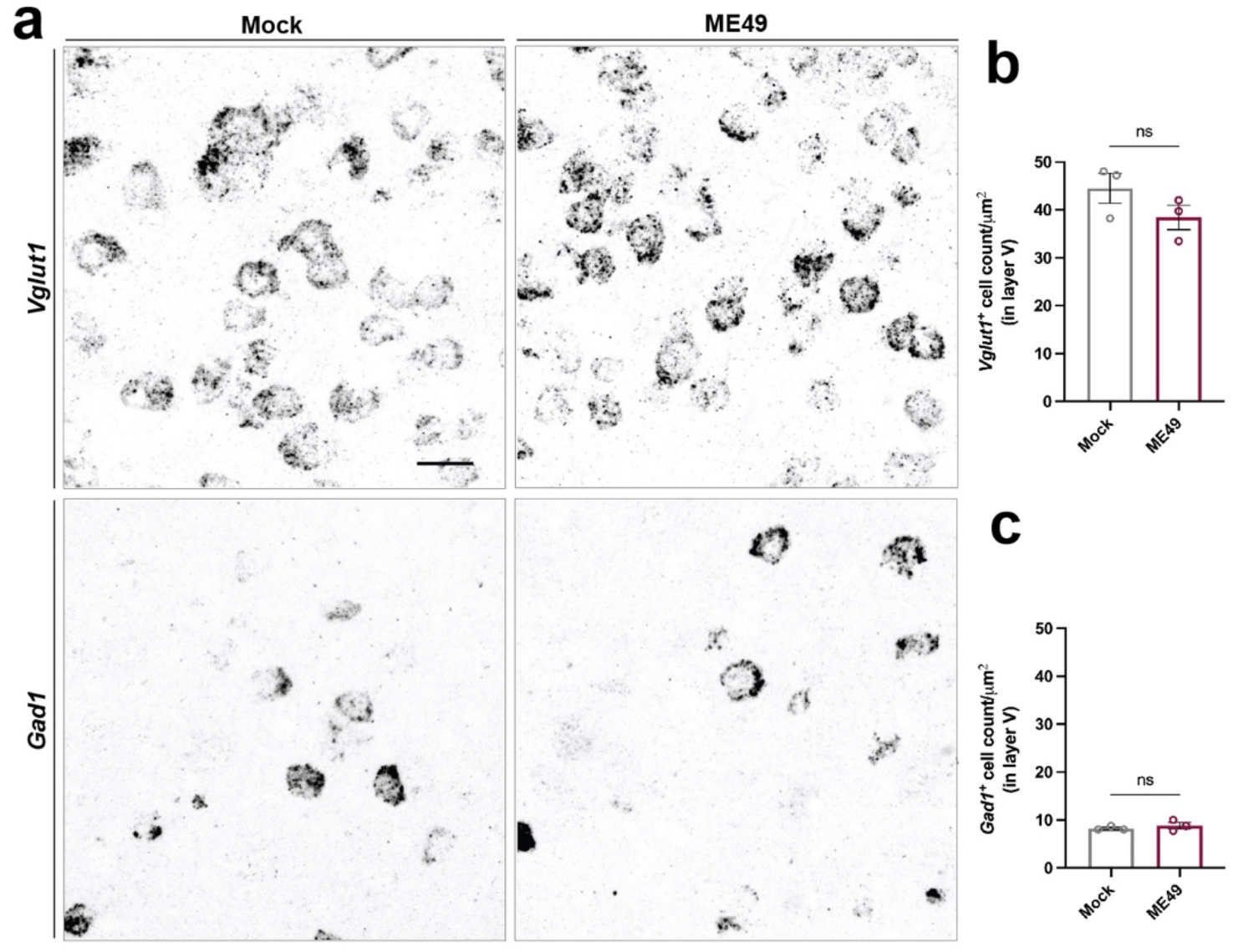
No excitatory or inhibitory neuron loss occurs at time of perisomatic synapse loss. (a) ISH for *Vglut1* and *Gad1* in layer V of mock- and ME49-infected neocortex reveals no significant decrease in the number of excitatory or inhibitory cells following infection. (b) Quantification of *Vglut1^+^* cell count in layer V of mock- and ME49-infected cortex. Each data point represents one biological replicate and bars depict mean ± SEM. No significant difference (ns) indicates *P* > 0.05 by Student’s *t*-test (n=3 mice per condition). (c) Quantification of *Gad1^+^* cell count in layer V of mock- and ME49-infected cortex. Each data point represents one biological replicate and bars depict mean ± SEM. No significant difference (ns) indicates *P* > 0.05 by Student’s *t*-test (n=3 mice per condition). Scale bar in A: 20 μm

**Supplemental Figure 2.**
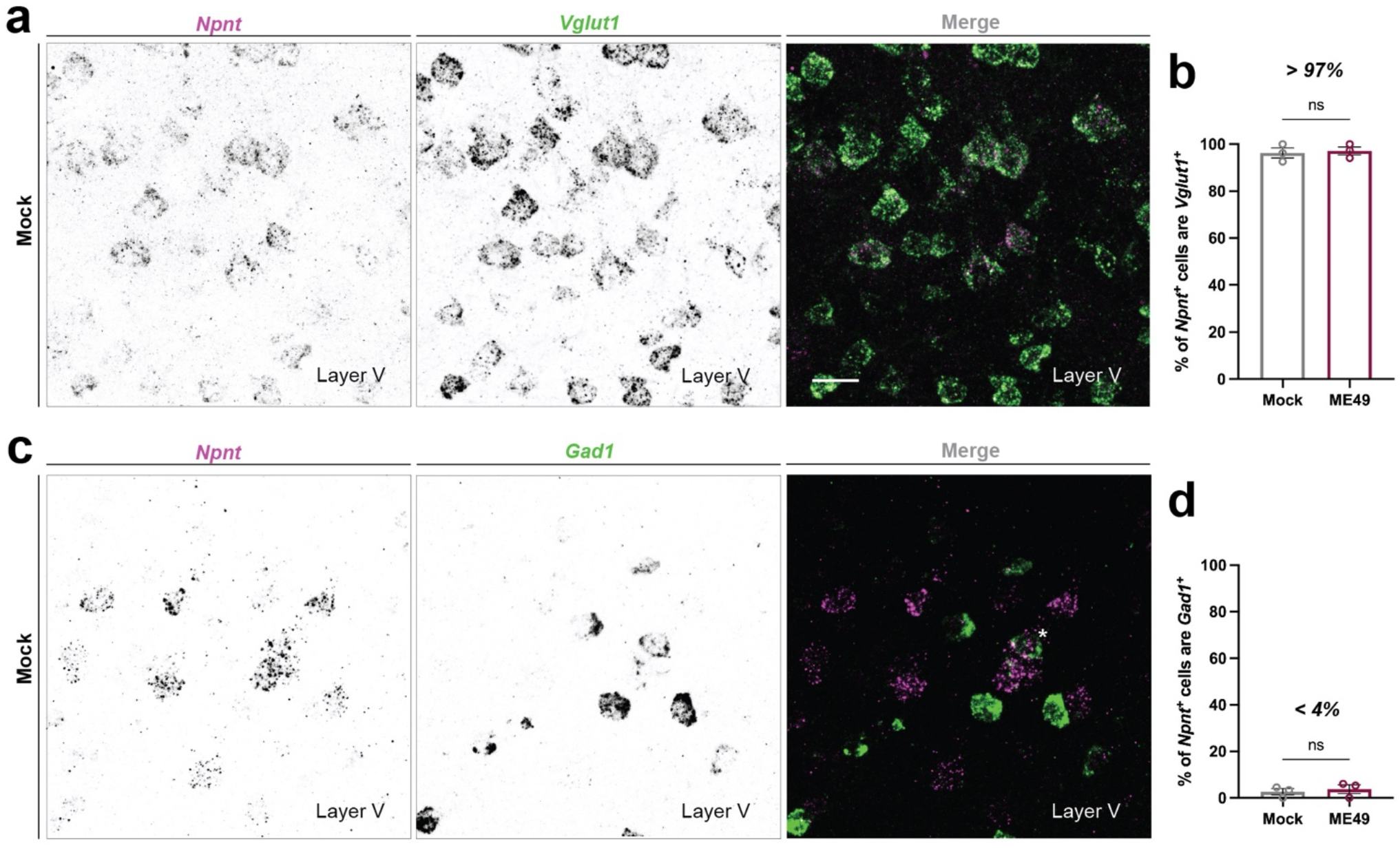
NPNT is a faithful marker of excitatory cells in cortical layer V. (a) ISH for *Vglut1* and *Npnt* in wildtype mouse neocortex shows most excitatory cells in layer V express *Npnt* mRNA. (b) Quantification of the percentage of *Npnt+* cells in layer V that express *Vglut1* mRNA in mock- and ME49-infected cortical layer V. Each data point represents one biological replicate and bars depict ± SEM. No significant difference (ns) indicates *P* > 0.05 by Student’s *t*-test (n=3 mice per condition). (c) ISH for *Gad1* and *Npnt* shows very few inhibitory cells in layer V express *Npnt* mRNA. Each data point represents one biological replicate and bars depict ± SEM. No significant difference (ns) indicates *P* > 0.05 by Student’s *t*-test (n=3 mice per condition). (d) Quantification of the percentage of *Npnt+* cells in layer V that express *Gad1* mRNA in mock- and ME49-infected cortical layer V. Scale bars in A and C: 20 μm

**Supplemental Figure 3.**
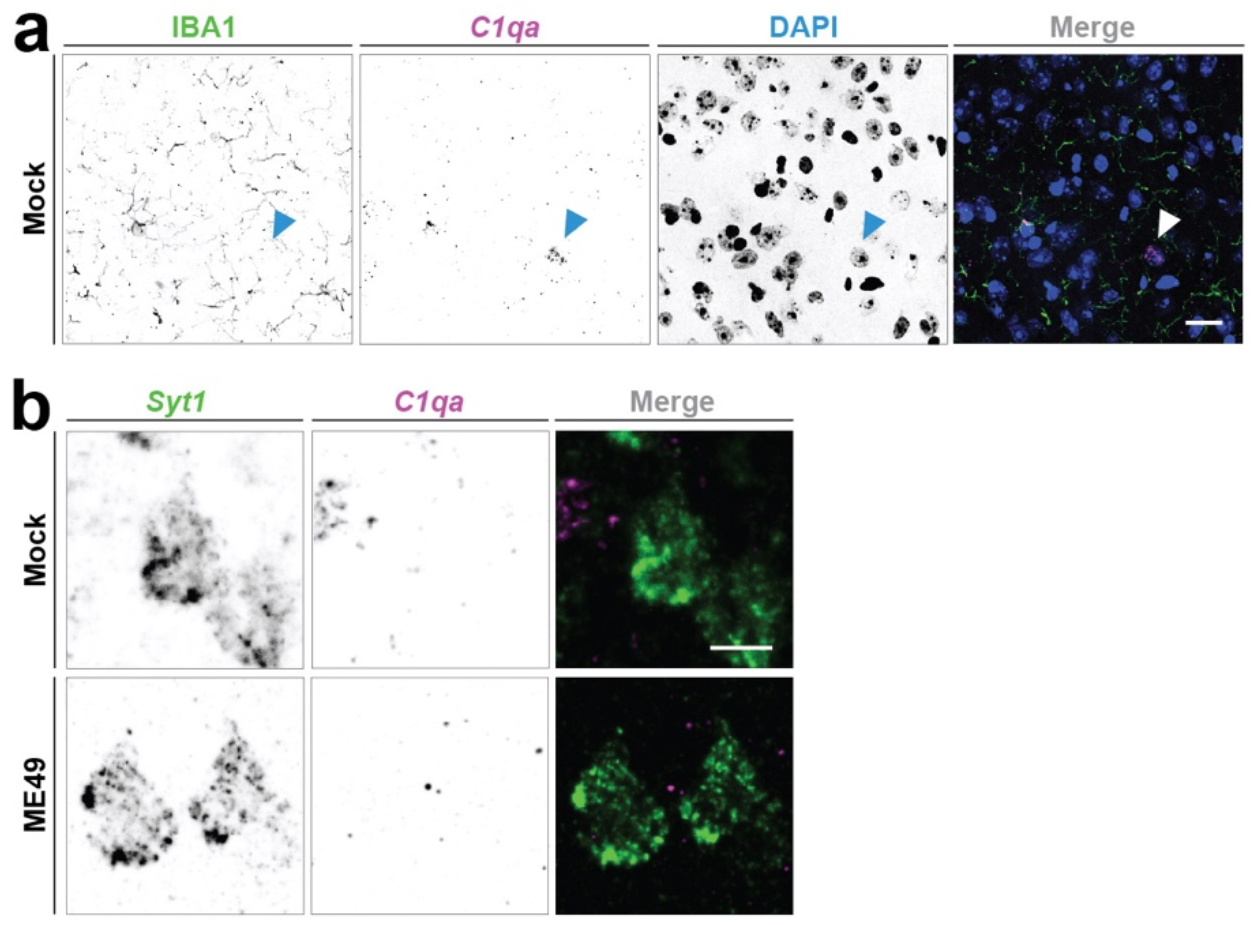
*C1qa* is not expressed by neurons. (a) ISH for *C1qa* and IHC for IBA1 and DAPI revealed *C1qa* mRNA expression by non-IBA1^+^ cell in mock-infected layer V. (b) ISH for *Syt1* and *C1qa* showed no neuronal expression of *C1qa* mRNA in both mock- and ME49-infected layer V. Scale bars in A: 20 μm and in B: 10 μm

**Supplemental Figure 4.**
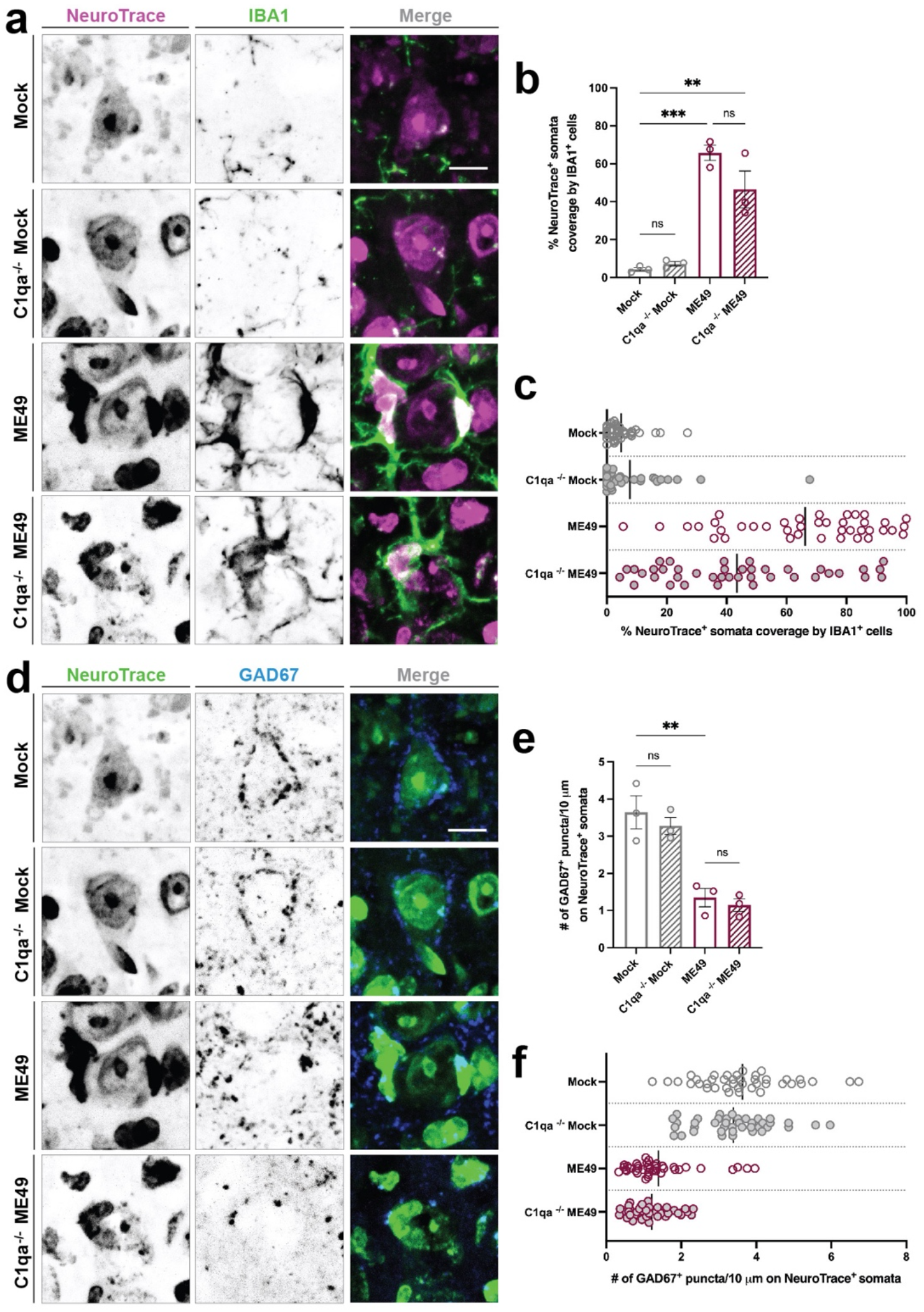
Inhibitory perisomatic synapse loss following *T. gondii* infection is not mediated by the classical pathway of complement activation. (a) IHC for IBA1 shows no reduction in microglial ensheathment of NeuroTrace^+^ neurons in C1qa^-/-^ ME49-infected cortex. (b) Quantification of percentage of NeuroTrace^+^ soma coverage by IBA1^+^ cells in cortical layer V in C1qa^+/+^ mock-, C1qa^-/-^ mock, C1qa^+/+^ ME49-. and C1qa^-/-^ ME49-infected brains. Each data point represents one biological replicate and bars depict mean ± SEM. Asterisks (**) indicate *P* < 0.01, (***) indicate *P* < 0.001, and (ns) *P* > 0.05 by one-way ANOVA with Sidak’s multiple comparison correction (n=3 mice per condition). (c) Distribution plot of individual NeuroTrace^+^ somata analyzed and pooled from biological replicates in (B). Bar represents mean percent somata coverage by IBA1^+^ cells. (d) IHC for GAD67 and NeuroTrace shows no reduction in inhibitory perisomatic synapse loss in C1qa^-/-^ ME49 infection compared to C1qa^+/+^ ME49-infection. (e) Quantification of GAD67^+^ synapses on NeuroTrace^+^ somata in cortical layer V in C1qa^+/+^ mock-, C1qa^-/-^ mock, C1qa^+/+^ ME49-. and C1qa^-/-^ ME49-infected brains. Each data point represents one biological replicate and bars depict mean ± SEM. Asterisks (**) indicate *P* < 0.01 and (ns) *P* > 0.05 by one-way ANOVA with Sidak’s multiple comparison correction (n=3 mice per condition). (f) Distribution plot of individual NeuroTrace^*+*^ cells analyzed and pooled from images in (E). Bar represents mean number of GAD67^+^ synapses on NeuroTrace^+^ somata. Scale bars in A and D: 10 μm

## MATERIALS AND METHODS

### Key resources table

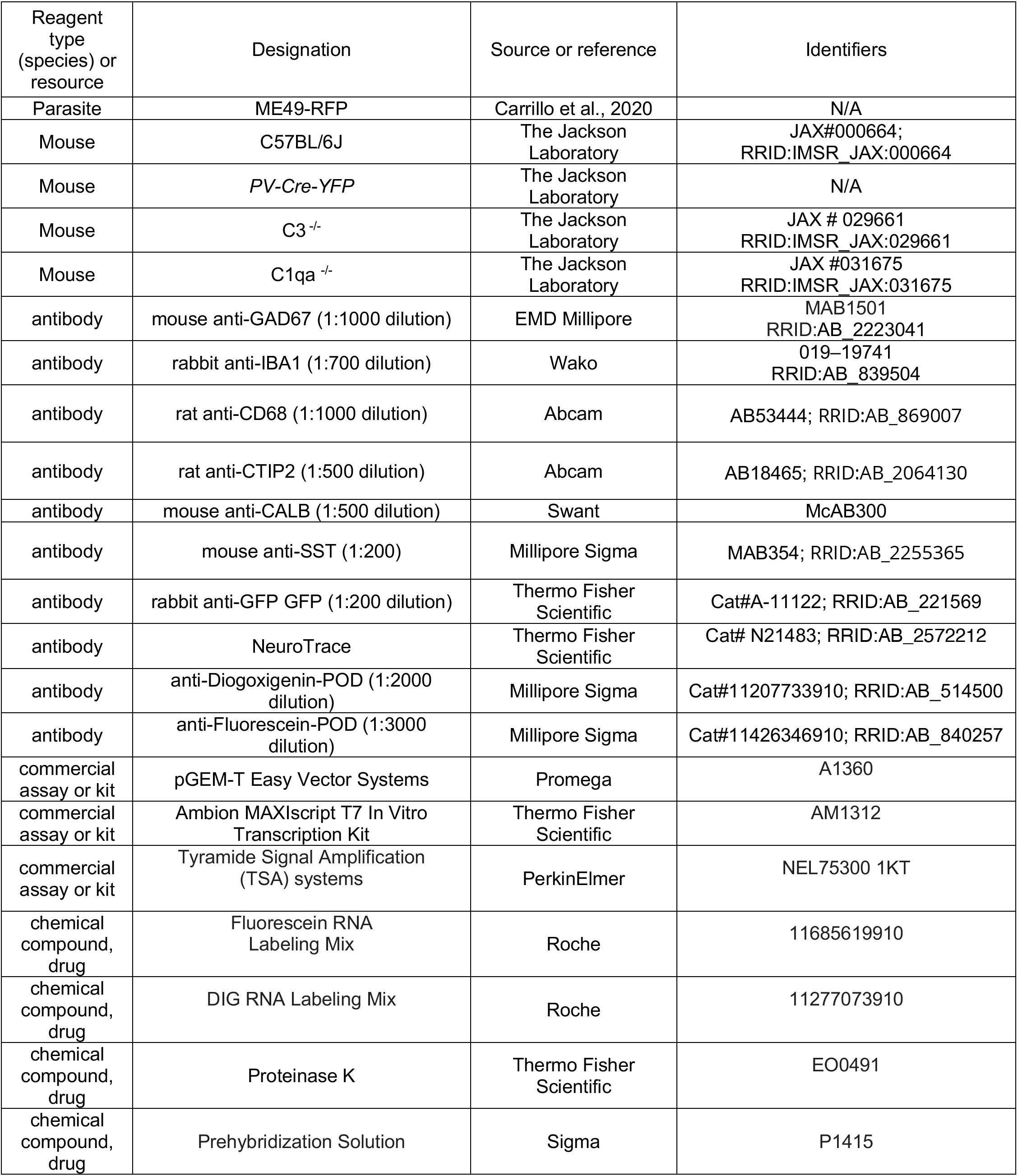
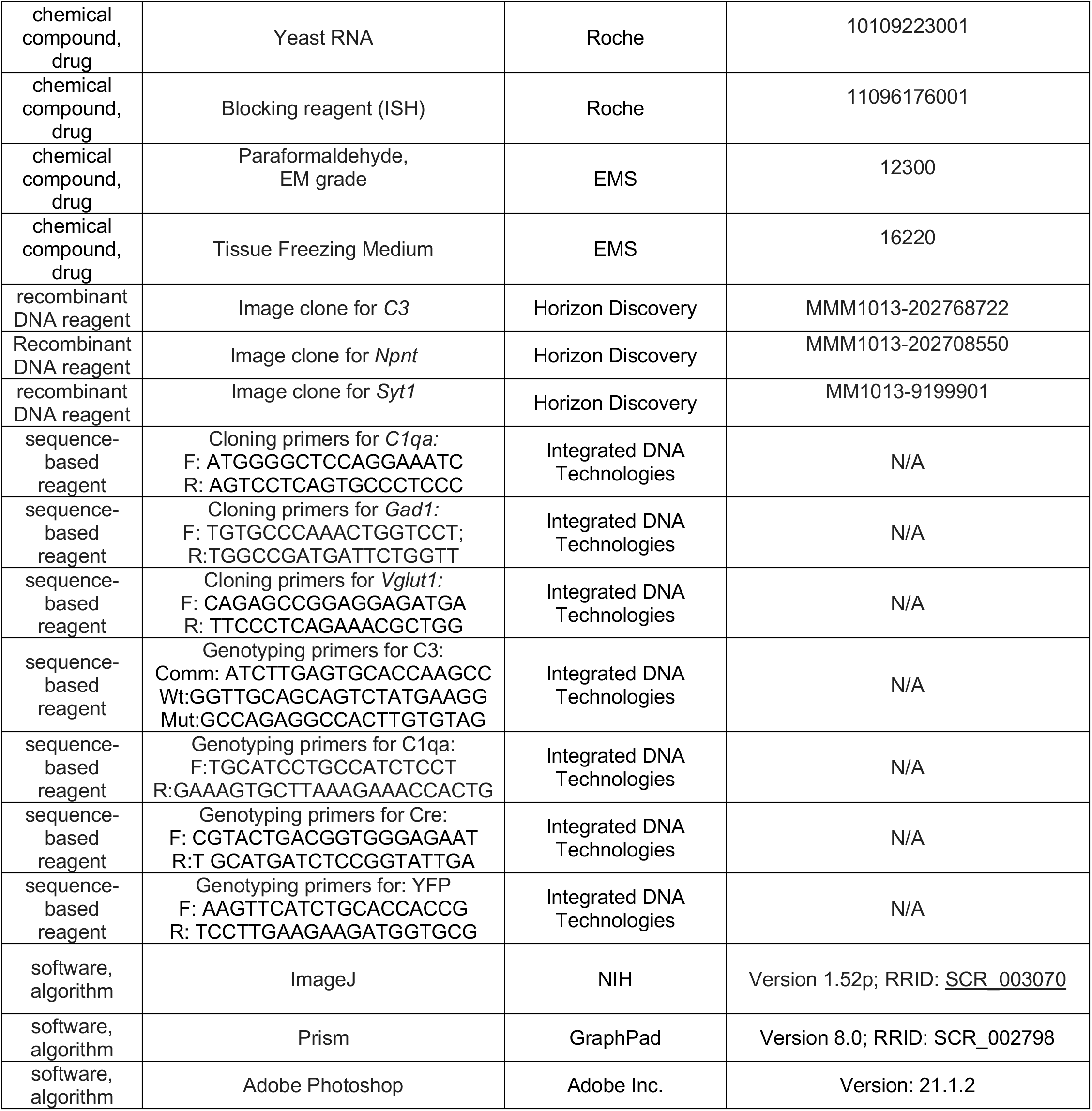

### Animals

C57/BL6 mice C3^-/-^ mice (JAX # 029661, genetic background: C57BL/6J), C1qa^-/-^ mice (JAX #031675, genetic background: C57BL/6J) were obtained from Jackson Laboratory. *PV-Cre-YFP* mice were generated from crossing *Parv-Cre*^KI/KI^ mice (JAX #008069, genetic background: C57BL/6J) with *Thy1-stop-yfp15* mice (JAX# 005630, genetic background: C57BL/6J) for several generations. All mice were genotyped before infection and experimentation (see key resource table for genotyping primer sequences). Both sexes were used for all experiments. Prior to infection, mice were housed in the same ABSL-1 temperature-controlled room with a 12 hr dark/light cycle and *ad libitum* access to food and water. Upon infection, mice were moved to an ABSL-2 room with similar temperature-control, 12 hr dark/light cycle, and they continued to have *ad libitum* access to food and water, supplemented with wet food. Experiments were performed in compliance with National Institute of Health (NIH) guidelines and protocols were approved by the Institutional Animal Care and Use Committee at Virginia Tech.

### Parasite infections

As previously described, age-matched male and female mice (~8 weeks of age) were infected with the Type II ME49-RFP strain of *T. gondii* (Brooks et al., 2015, Carrillo et al., 2020). The total number of *T. gondii* cysts within 20μl of infected brain homogenate was determined by fluorescence microscopy. Mice were interperitoneally infected with 2 ME49-RFP cysts in PBS. For time course experiments, brains were collected after 7 days, 12 days, 21 days, and 30 days of infection. For long-term infection, brains were collected 30 days following infection.

### Tissue Preparation

Tissue was prepared as previously described in Su et al., 2010. Briefly, an intraperitoneal injection of tribromoethanol at a concentration of 12.5 ug/ml was used to anesthetize mice. Mice were perfused with 10 mL of DEPC-treated 1x PBS followed by DEPC-treated 4% PFA pH 7.4. Brains were kept overnight at 4°C in 4% DEPC-PFA and then transferred to 30% DEPC-sucrose for at least 4 days. Fixed brains were embedded in Tissue Freezing Medium (Electron Microscopy Sciences), frozen overnight, and coronally sectioned on a Leica CM1850 cryostat at 16μm thickness. Prepared tissue slides were kept in −20°C for later IHC and ISH experiments.

### Immunohistochemistry (IHC)

Tissue slides were air-dried at room temperature for 15 minutes. Slides were incubated in IHC blocking buffer (2.5% bovine serum albumin, 5% normal goat serum, 0.1% Triton-X in PBS) for 1 hour. Primary antibodies were diluted in blocking buffer at the concentration described in the key resource table. Tissue slides were incubated in diluted primary antibody at 4°C for a minimum of 15 hrs. After removal of primary antibody, slides were washed in 1x PBS for an hour, followed by incubation with fluorophore-conjugated secondary antibodies diluted IHC blocking buffer at 1:1000 for 1 hour room temperature. Lastly, after several washes in 1x PBS, tissue sections were either stained with 4’,6-diamidino-2-phenylindole (DAPI) or with NeuroTrace 640/660. NeuroTrace staining was performed as follows: First, slides were incubated in 0.1% Triton-X in PBS for 10 minutes, washed in 1x PBS twice for 5 minutes each, incubated in NeuroTrace (diluted 1:250 in PBS) for 35 minutes in room temperature, washed in 0.1% Triton-X in PBS for 10 minutes, and two final washes in 1x PBS.

### Riboprobe production

pCMV-SPORT6 plasmids carrying *C3* (cat # 5134713), *Syt1* (cat # 5363062), and *Npnt* (cat # 5149204) were obtained from Horizon Discovery (previously GE Dharmacon). *Gad1* cDNA, *C1qa* cDNA, and *Vglut1* cDNA were generated and amplified as previously described in Monavarfeshani et al., 2018. Plasmids and primers are listed in the key resource table. All riboprobes were generated as described in Monavarfeshani et al., 2018.

### In situ hybridization (ISH)

ISH was performed using the in-house generated riboprobes on fixed tissue, prepared with DEPC-treated reagents as described above. Tissue sections on slides were fixed in 4% DEPC-PFA for 10 minutes at room temperature, washed with 1x DEPC-PBS, and incubated with proteinase K solution (1ug/mL in 50 mM Tris pH 7.5, 5 mM EDTA) for 10 minutes at room temperature. Slides were then washed with 1x DEPC-PBS, incubated in 4% DEPC-PFA for 5 minutes at room temperature, and incubated in acetylation solution (0.25% acetic anhydride, 20 mM hydrochloric acid and 1.33% triethanolamine) for 10 minutes at room temperature. To permeabilize the tissue, slides were incubated in 1% triton in DEPC-PBS for 30 minutes room temperature. To block endogenous peroxidase, slides were incubated in 0.3% hydrogen peroxide in DEPC-PBS for 30 minutes at room temperature, followed by DEPC-PBS washes. Tissue sections were equilibrated in hybridization solution (40 mL of prehybridization solution, 1.6 mL of 5 mg/mL, and 25 mg Roche yeast RNA) for 1 hour at room temperature. Riboprobes were denatured for 10 minutes at 80°C, applied to tissue sections, cover slipped, and incubated overnight at 65°C (probes were denatured for 10 minutes at 80°C). On day 2, coverslips were removed by slide incubation in 2x saline-sodium citrate solution for 5 min in 65°C, and slides underwent several 45 minute-long washes at 65°C, with a final wash at room temperature, followed by rinsing with 0.2x saline-sodium citrate solution in tris-buffered saline (TBS) at room temperature. Slides were incubated in ISH blocking buffer (10% lamb serum 0.2% Roche blocking reagent in TBS) for 1 hour at room temperature and then incubated overnight at 4°C in HRP-conjugated anti-DIG or anti-FL antibodies (in key resource table). On day three, riboprobes were identified using a Tyramide Signal Amplification (TSA) system (in key resource table). For double riboprobe ISH, the first riboprobe antibody was quenched by incubation in 2% hydrogen peroxide in TBS for 1 hour and 15 minutes at room temperature. After washing with TBS, slides were incubated in ISH blocking buffer for 1 hour at room temperature and incubated overnight at 4°C in the second HRP-conjugated anti-DIG or anti-FL antibody. On day four, the second riboprobe was identified with the TSA system and then washed with TBS. For IHC following ISH, slides were incubated in IHC blocking buffer and IHC was performed as described above. IHC antibody dilutions following ISH were as follows: rabbit anti-GFP dilution: 1:200, rabbit anti-IBA1 dilution: 1:600).

### Imaging

Images were acquired with a Zeiss LSM700 confocal microscope. Representative images shown in figures are of a maximum intensity projection, unless otherwise noted. Analyses were performed with single optical section images.

### Density of cells (*Vglut1*^+^, *Gad1*^+^, *PV-Cre-YFP*^+^ *IBA1*^+^)

For assessment of cell numbers in cortical layer V, images were obtained using 40x magnification (20x 2 zoom). Counts were performed manually. The “Count Tool” function in Adobe Photoshop (Adobe Inc., version: 21.1.2) was used to keep track of cell counts and colocalized counts. Layer IV – VI cell counts were obtained from 20x images. Cell counts are normalized by the ROI area described as μm^2^.

### Cell type ensheathment analysis

Excitatory and inhibitory cell types were labeled either by immunohistochemistry (CTIP2, SST, CALB) or by use of a transgenic reporter *(PV-Cre-YFP;* details in ‘Animals’ section). Tissue was co-stained with NeuroTrace for visualization of the entire neuronal soma (except for *PV-Cre-YFP).* Images were taken at 40x magnification to capture only cortical layer V at 1μm intervals. Cells of interest were identified by colocalization of cell type marker with NeuroTrace, and the % somata coverage by IBA1^+^ cell was calculated by length of IBA1^+^ cell or process contact neuronal soma (μm) / neuronal soma perimeter (μm) for each optical section where the cell’s NeuroTrace signal was visible. Length and perimeter measurements were taken with ImageJ. % soma coverage calculations on individual optical sections were averaged to obtain % soma coverage for each cell. Data is shown as quantification of biological replicates (mean of all cells from each animal) and plotted as individual cell data points to show distribution across all animals.

### Inhibitory perisomatic synapse analysis

Cells were labeled transgenically *(PV-Cre-YFP)* or by NeuroTrace. Inhibitory perisomatic synapses were identified with GAD67 immunolabeling. Images were taken at 40x magnification to capture only cortical layer V at 0.5μm intervals. The number of GAD67^+^ puncta contacting the perimeter of NeuroTrace-labeled neuronal soma were counted manually and normalized to the neuronal soma perimeter (measured by ImageJ) at each optical section. Calculations from optical sections were averaged to obtain a single calculation of the number of GAD67^+^ puncta / 10μm perimeter of neuronal soma. Data is shown as quantification of biological replicates (mean of all cells from each animal) and plotted as individual cell data points to show distribution across all animals.

### Colocalization analysis

All ISH colocalization *(Syt1* and *C3, Syt1* and *C1qa, Npnt* and *C3*, *Gad1* and *C3*, *Npnt* and *Vglut1, Npnt* and *Gad1)* was determined by single optical image sections. The “Count Tool” function in Adobe Photoshop was used to keep track of cell counts and colocalized counts.

### IBA1 masked analysis (*C1qa* and CD68)

IBA1^+^ cells were masked using the ‘threshold’ and ‘analyze particles’ functions in ImageJ. The masks were superimposed to either the CD68 channel, or the *C1qa* channel and the mean signal intensity was calculated within the masks, and outside the masks (by inverting the ROI).

### Expression analysis (% of IBA1^+^ cells express *C1qa*, % of *Npnt*^+^ cells express *Vglut1* or *Gad1*)

Quantifications for the percentage of IBA1^+^ cells that express *C1q* were determined by maximum intensity projection images (and confirmed by single optical sections). The number of IBA1^+^ cells with colocalized *C1q* signal was divided by the total number of IBA1^+^ cells per image area (μm^2^). For % of *Npnt^+^* cells that express *Vglut1*^+^ or *Gad1,* the number of *Vglut1\** cells (or the number of *Gad1*^+^ cells) that colocalized with *Npnt*^+^ cells were divided by the total number of *Npnt*^+^ cells per image area (μm^2^). All cells were counted manually and the “Count Tool” function in Adobe Photoshop was used to keep track of cell counts and colocalized counts.

### Quantifications and statistics

All analyses were performed with 3-4 biological replicates per genotype and condition. Both sexes were used in the quantification. No data or animals were excluded. No sex-specific differences were observed with the number of animals used. Statistical analyses (Student’s *t*-test or one-way ANOVA with Sidak’s correction for multiple comparisons as indicated in figure legends) were performed using GraphPad Prism (version 8.0; RRID: SCR_002798). *P* < 0.5 values were considered to be significantly different (*P* values included in figure legends). Data points represent biological replicates or individual cells analyzed as described in figure legends and are plotted as ± SEM or mean, respectively.

